# Generalisable signatures of anaesthesia in the large-scale functional organisation of the marmoset brain

**DOI:** 10.64898/2026.06.06.730576

**Authors:** Sebastian Dohnany, Katarina Jerotic, Davide Orsenigo, Elisa Serra, Hana Ali, Jonathan Buhler, Zhen-Qi Liu, Kanako Muta, Junichi Hata, Hideyuki Okano, Gustavo Deco, Morten L. Kringelbach, Andrea I. Luppi

## Abstract

How the activity and connectivity of the brain support consciousness remains a central question in neuroscience. Recent progress driven by the use of functional MRI has seen growing recognition that large-scale distributed functional organisation of the human and non-human primate brain are systematically and consistently reshaped by anaesthetic-induced unconsciousness, across anaesthetics and across human and macaque. Here, we generalise these results to a different primate species that is gaining traction as model organism in neuroscience, the marmoset (*Callithrix jacchus*). We also generalise results to an additional anaesthetic, isoflurane, which we compare with propofol and sevoflurane. We report that under anaesthesia with propofol, sevoflurane, or isoflurane, distributed brain activity from functional MRI is increasingly constrained by the underlying structural connectivity across scales. Anaesthesia also induces a collapse of the principal gradient and intrinsic functional geometry of the marmoset brain, coinciding with a breakdown of hierarchical integration. Altogether, the present results indicate generalisable signatures of anaesthesia in the large-scale organisation of the primate brain.

## INTRODUCTION

How the activity and connectivity of the brain underpin various states of consciousness remains an open question. Historically, the focus has mainly been on the specific role of brain regions or the functional interactions between them, in some cases widening the view to consider how spatially and functionally distant networks reorganization coincide with altered states of consciousness [1]. Recent neuroimaging studies, however, indicate that this perspective only partially captures the underlying complexity [1, 2]. Both pathological and pharmacological disruptions of consciousness are associated with a whole-brain reorganization that unfolds along both functional and anatomical hierarchies of brain organization [3–5]. Such large-scale organization and structure–function relationships can be observed through the lens of functional gradients [4, 6–11] and connectome harmonics (i.e., the eigenmodes of the structural connectome) [12–16]. For instance, recent studies have demonstrated that cortical gradients of functional connectivity can serve as reliable markers of loss of consciousness. In humans, degradation of these gradients correlates with a variety of pathological and pharmacologically induced states [5], and similar patterns have now been observed in macaque models across multiple anaesthetics [4]. Similarly, connectome harmonics — which offer a quantitative framework for assessing the correspondence between structural and functional brain connectivity across scales (from individual regions to the entire cortex) [12] — have been shown to effectively characterize altered states of consciousness from a neural dynamic perspective [3, 4, 13]. Notably, these measures reveal a significant increase in structure-function coupling during anesthesia and in disorders of consciousness, a finding that holds across both human subjects and macaque models [2].

Previous work examined the effects of sevoflurane and propofol – anaesthetics with distinct molecular mechanisms – on the redistribution of large-scale patterns of cortical functional organisation in macaque brains [4]. These changes were examined using three key markers: harmonic energy [3, 4, 12], gradient range [5], and hierarchical integration [4, 17].

Marmosets (*Callithrix jacchus*) are an invaluable non-human primate (NHP) model for studying various neurobiological and functional properties, particularly in the context of anaesthesia and cortical organisation. Their importance stems from two key factors: their strong translational value for human neuroscience research, and their amenability to genetic manipulation. As primates, marmosets share crucial similarities with humans in cortical organisation, functional architecture, and behavioural traits, making them an ideal model for studying higher-order neural processing [18–21]. Furthermore, the effects of general anaesthesia on the interactions between cortical regions are highly conserved across mammals [3, 4, 22, 23], reinforcing the predictive power of marmosets for human anaesthesia research.

Beyond their translational relevance, marmosets provide a unique opportunity to extend anaesthesia research to a different type of primate brain architecture. Most existing studies on anaesthesia in NHPs have focused on macaques, which possess highly folded (gyrencephalic) cortices, like humans. In contrast, marmosets possess a smooth (lissencephalic) cortex, like rodents [24, 25]. This makes marmosets a critical translational bridge between rodents - lissencephalic, but commonly used in preclinical trials - and macaques, whose more complex and highly folded brains are closer to human neuroanatomy, but less amenable to experimental accessibility [26]. Marmosets’ enhanced susceptibility to genetic modification explains the heightened interest in this species for neuropsychiatric modelling [21, 27, 28].

Despite an increasing interest in using marmosets in neuroscience, in particular to study functional and structural connectivity [29], and promising results in using marmosets to study structure-function relationship using eigenmode decomposition [30] and functional gradients [31], investigations of the effects of anaesthesia in this model organism remain sparse. In particular, to date, no studies have yet investigated how structure-function relationships and functional gradients in the marmoset brain are altered under anaesthesia.

In the current paper, we examine changes in recently developed markers of consciousness (functional gradients and connectome harmonics) following anaesthesia administration in a previously published marmoset resting state fMRI dataset [24]. We aim to replicate changes induced by sevoflurane and propofol administration within the marmoset brain, as previously observed with macaque data. Further, we explore the effects of isoflurane, an anaesthetic whose effects on such redistributions of cortical functional organisation and structure-function interactions have not previously been examined in either species. Thus, our goal is to generalise distributed cortical markers of consciousness to a different species (marmoset) and a different anaesthetic (isoflurane).

## RESULTS

### Structural Harmonics

We examine the interdependence of functional connectivity on underlying anatomical structure following anaesthesia in marmosets by applying harmonic mode decomposition to fMRI data. Harmonic mode decomposition is a mathematical framework that decomposes brain activity into multi-scale patterns of co-activation, based on how functional signals propagate over the brain’s structural connectome. Each resulting component of activity is represented by a harmonic mode, which varies in spatial granularity and is ordered by increasing frequency and complexity as a function of the corresponding eigenmodes (Figure 1A). Low-frequency harmonic modes correspond to large-scale activity that is highly constrained by the underlying structural pathways (e.g., activity spanning both hemispheres). In contrast, high-frequency harmonic modes capture more localized, fine-grained activity, which is less dependent on structural connectivity, allowing for more functionally flexible interactions (Figure 1B-C). The relative contribution of high-vs. low-frequency harmonic modes in the fMRI data provides insight into structure-function coupling. High harmonic energy indicates a stronger dependence of functional activity on structural connectivity, reflecting a more constrained and less flexible functional organization.

**Figure 1.**
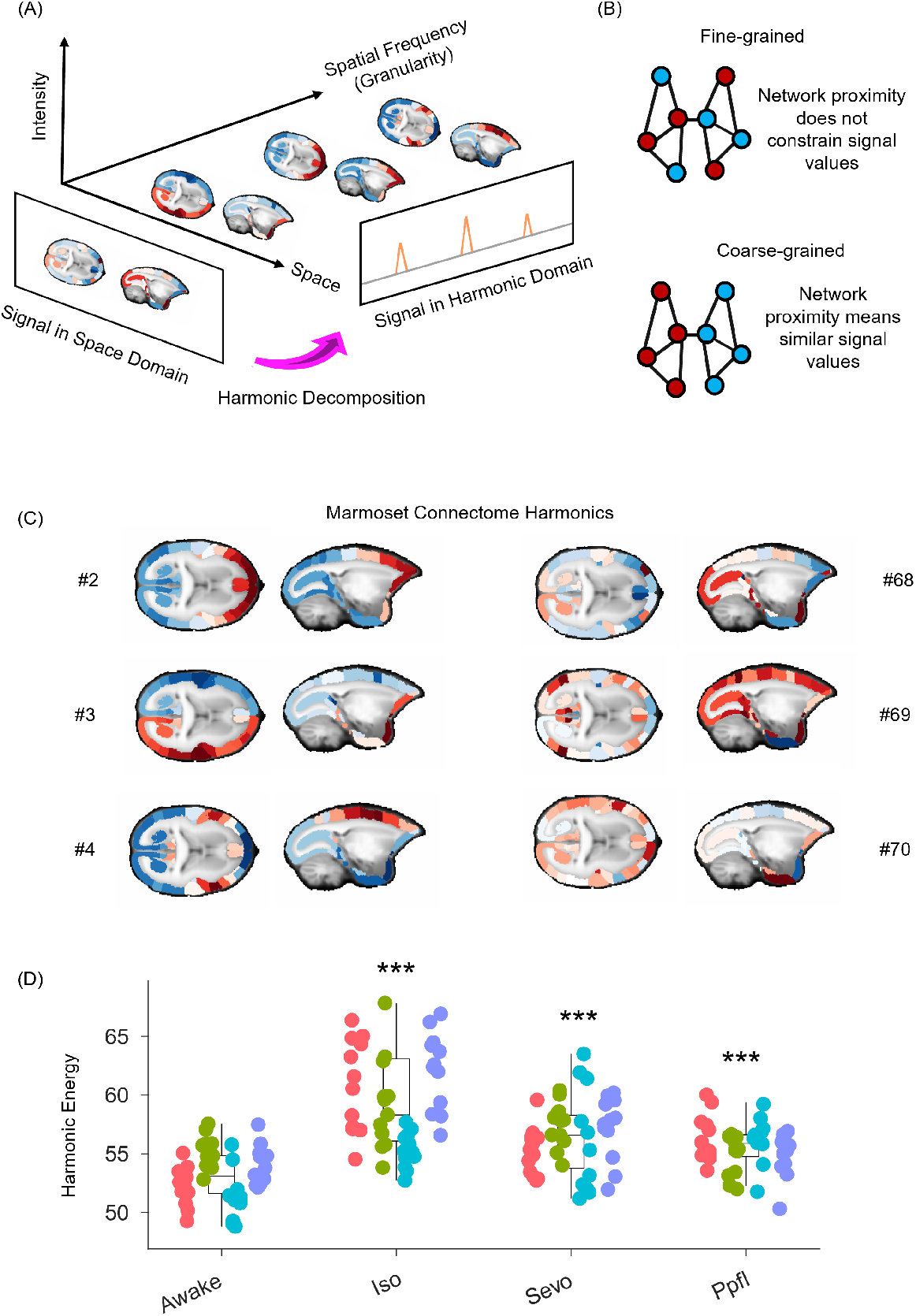
Anaesthesia increases structure-function coupling in the marmoset brain. **A)** Harmonic decomposition of the connectome takes a spatial signal — the fMRI BOLD activation recorded at specific cortical locations — and expresses it in terms of harmonic modes defined by the brain’s structural connectivity. This yields a set of basis functions that capture distributed activity patterns across the entire brain at various spatial scales (granularity). These range from broad, smooth gradients along principal anatomical axes (notably left–right and anterior–posterior) to progressively finer, detailed patterns. Note that here, frequency is not about time, but about spatial scale. **B)** Low-frequency (coarse-grained) harmonics correspond to spatial patterns closely related to the underlying structural connectome. Strongly connected nodes exhibit similar activation (indicated by colour). High-frequency (fine-grained) patterns are characterized by divergence between the spatial organisation of the functional signal and the underlying network structure. Nodes closely connected in the structural network might exhibit different functional signals. **C)** Represented on the marmoset brain are the first three and the last three non-uniform harmonic modes of its structural connectome. The initial, low-frequency modes reveal broad, large-scale patterns, while the latter, high-frequency modes capture finer, detailed patterns. The total number of harmonics—70 in this case—matches the number of brain regions in the connectome, establishing the maximum achievable resolution. **D)** The energy (contribution) of harmonic modes is significantly increased in the marmosets brain under deep anaesthesia, whether induced by isoflurane, sevoflurane, or propofol. \**p* < 0.05; **, *p* < 0.01; \*\*\**p* < 0.001 from linear mixed effects modelling (two-sided, FDR-corrected), compared against Awake condition. N=12 runs for 4 animals for each conditions (e.g. Awake, Isoflurane, Sevoflurane and Propofol). Box plots: central line, median; box limits, upper and lower quartiles; whiskers, 1.5× interquartile range; Data points of the same color correspond to the same animal. Statistical comparisons from linear mixed effects modeling (\*\*\**p* < 0.001, two-sided, FDR-corrected) are made relative to the awake condition; see Table S1 information for full statistical results.

Here we observed significant (FDR-corrected) harmonic energy increases following anaesthesia administration, denoting an increase in structure-function coupling, or increased structural constraints on observed brain activity, as compared to the waking state. As shown in Figure 1D, this significant increase is observed across all three anaesthesia conditions (isoflurane: *β*(5.036, 7.312) = 6.174, *p* < 0.0001; sevoflurane: *β*(2.230, 4.158) = 3.194, *p* < 0.0001; propofol: *β*(1.564, 3.225) = 2.394, *p* < 0.0001).

### Gradients of Functional Geometry

We examine the distribution of brain function by exploring the principal functional gradient. Functional gradients can be thought of as nonlinear variations of principal components analysis [6, 7]. Like PCA, functional gradients outline patterns most representative of the underlying brain activity through eigenmodes and eigen-vectors, aligning with the dimensions of the functional connectivity’s spatial variation. Functional proximity between regions in the brain is denoted with more similar gradient values (and therefore less variation), whereas regions whose functional connectivity with the rest of the brain show the greatest difference will make up the extremes of the gradient [6, 7] (Figure 2A).

**Figure 2.**
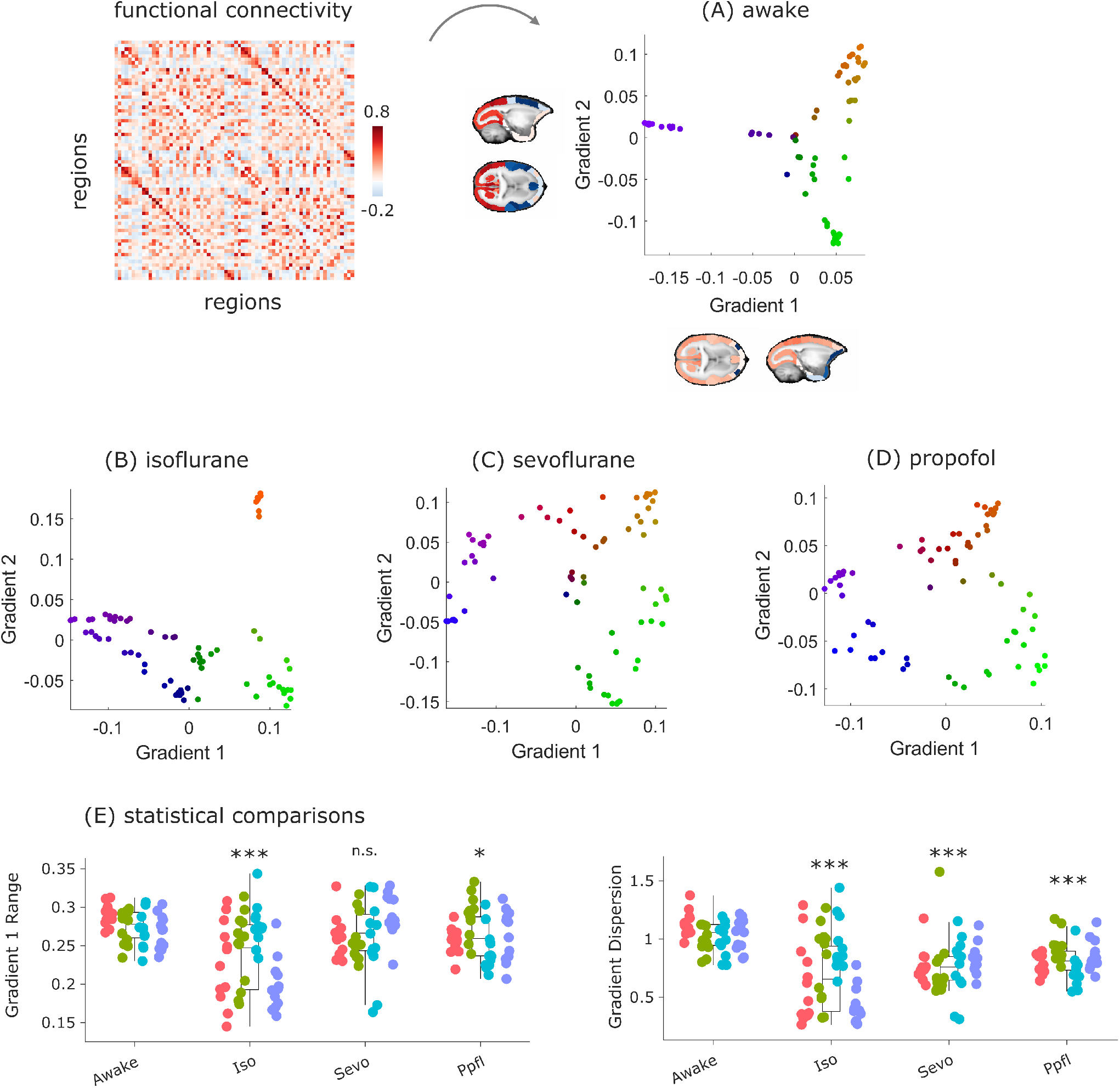
Anaesthesia collapses the intrinsic functional geometry of the marmoset brain. **A)** The nonlinear low-dimensional embedding of marmoset functional connectivity using the first two gradients, which were derived via diffusion graph embedding (see “Methods”) from the group-averaged FC matrix in the awake condition. These gradients are also projected onto the marmoset brain, with colors indicating each region’s position along the gradients. **B**,**C**,**D)** Scatter plots display the first two principal gradients derived from the group-averaged FC matrix for each anesthetized condition (isoflurane, sevoflurane, and propofol). The dots’ colors indicate the extremes of the gradients: green and blue mark the endpoints of the first gradient, while red and blue mark those of the second gradient. **E)** The range of the principal gradient of marmoset functional connectivity across wakefulness and various anesthetic conditions, along with the gradient dispersion calculated from the first three gradients. Box plots: central line, median; box limits, upper and lower quartiles; whiskers, 1.5× interquartile range; Data points of the same color correspond to the same animal. Statistical comparisons from linear mixed effects modeling (\*\*\**p* < 0.001, two-sided, FDR-corrected) are made relative to the awake condition; see Tables S2 and S3 for full statistical results.

We start by exploring the gradient dispersion, which gives us a measure of the average Euclidean distance of each datapoint across the first three gradients compared to the centroid across the three. Results show significant reductions in measure of gradient dispersion, greater average Euclidean proximity, across all anaesthesia conditions (isoflurane: *β*(-0.438, -0.237) = -0.338, *p* < 0.0001; sevoflurane: *β*(-0.331, -0.190) = -0.260, *p* < 0.0001; propofol: *β*(-0.272, -0.161) = -0.216, *p* < 0.0001) (Figure 2B-E.

In humans, the principal functional gradient is thought to show the greatest spatial variation and has been suggested to reflect the separation between unimodal and multimodal brain regions: those sitting at either extremes of the cortical functional hierarchy [10]. Previously, we showed significant reductions in gradient range under deep propofol and ketamine in macaques, whereas no significant differences were observed in the case of sevoflurane [4]. In the current study, using linear mixed effects models, we show FDR-corrected significant reductions in gradient range under both isoflurane (*β*(-0.053, -0.024) = -0.038, *p* < 0.001) and propofol (*β*(-0.023, - 0.002) = -0.013, p = 0.029). No FDR-corrected significant difference, compared to the awake condition, is observed under sevoflurane (*β* = -0.009, p = 0.145) (Figure 2B-E; see Tables S2 and S3 for full statistical results).

### Hierarchical integration and segregation

Finally, we examine changes to the functional signal’s hierarchical integration and segregation following anaesthesia administration. Changes to the balance of integration and segregation dynamics across scales is a fundamental feature of many theories of consciousness [32–35]. Unlike many classic measures of integration and segregation which explore changes in integration and segregation at one scale [17, 36, 37], here we take a multi-scale approach, by considering all functional eigenmodes [38]. Each eigenmode extracted from functional connectivity delineates a unique pattern or group of regions which show cohesive activation patterns (same sign) or alternate activation patterns (opposite sign). Due to the nature of eigenmodes, progressively showing greater division into more and iteratively smaller modules, we can outline hierarchical modules based on the alignment in signs, or lack thereof, between regions across eigenmodes [4, 38]. Importantly, here we explore nested relationships between elements of the system across scales, as modules which are segregated at one hierarchical level may be integrated in the one before it [38].

We examine changes to hierarchical segregation under anaesthesia, by computing the cumulative weighted contribution of all eigenmodes, excluding the first (see Methods). In the current analyses, all anaesthetic conditions showed significant reductions in hierarchical segregation scores compared to the awake state, as observed through linear mixed effects model analyses (isoflurane: *β*(-0.032, -0.017) = -0.024, *p* < 0.0001; sevoflurane: *β*(-0.060, - 0.046) = -0.053, *p* < 0.0001; propofol: *β*(-0.030, -0.017) = -0.024, *p* < 0.0001) (Figure 3).

**Figure 3.**
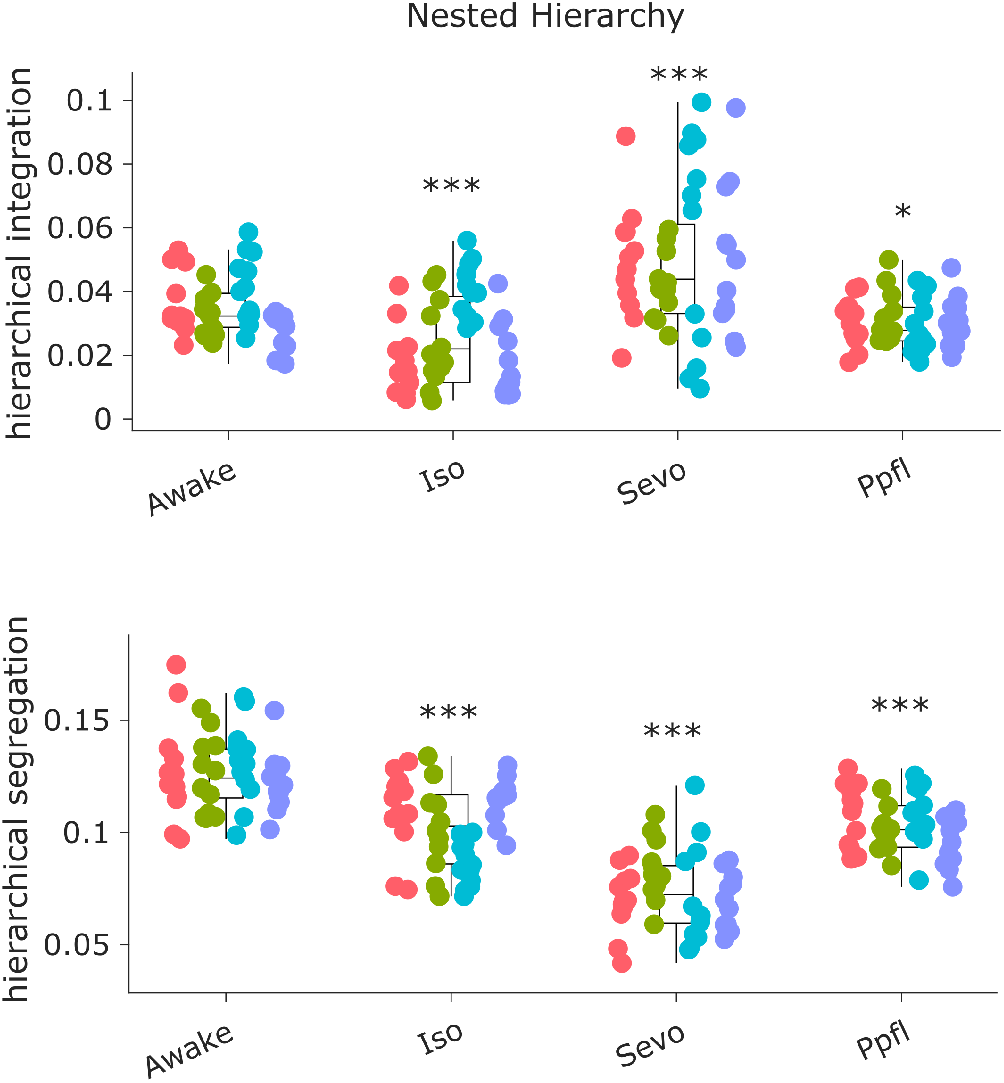
Anaesthesia reshapes nested hierarchies in the marmoset brain. **Top)** Eigenmode-based hierarchical integration is significantly diminished in the marmoset brain under anesthesia induced by isoflurane or propofol, while it is increased with sevoflurane. **Bottom)** Eigenmode-based hierarchical segregation is reduced across all anesthetic conditions. Statistical significance was determined using linear mixed effects modelling (two-sided, FDR-corrected; \**p* < 0.05, \*\*\**p* < 0.001, FDR-corrected) relative to the awake condition. Box plots: central line, median; box limits, upper and lower quartiles; whiskers, 1.5× interquartile range; within each panel, dots of the same colour are provided by the same animal. See Tables S4 and S5 for full statistical results.

Hierarchical integration is delineated by the contribution of the first eigenmode, as this eigenmode has the same sign across the whole cortex. Aligned with previous findings in macaque fMRI data, linear mixed effects model results show a significant reduction in hierarchical integration following administration of propofol (*β* (-0.008, -0.001) = -0.004, *p* = 0.017). The same significant decrease was observed following the administration of isoflurane (*β*(-0.014, -0.005) =-0.010, *p* < 0.0001). In the case of sevoflurane, the opposite trend was observed, namely a significant increase in hierarchical integration (*β*(0.008, 0.022) = 0.015, *p* < 0.0001), suggesting differential drug effects on hierarchical integration (Figure 3).

### Robustness analysis

Diffusion map embedding has a number of free parameters [6, 7]. In line with previous work, for our main analyses we used the default options from the BrainSpace toolbox [4, 5, 7]. In Supplementary Figures S1-S10 we show that our main results from gradient analysis of functional geometry are robust to specific parameter choices, including different sparsity levels of the graph; different values of the *α* parameter that controls the balance between local and global structure in the nonlinear embedding; and whether the gradients are aligned to the same reference gradients from awake animals using Procrustes alignment, or determined in a data-driven manner for each individual [4, 7]. As with our main analysis, we find that gradient dispersion (which captures geometric changes across the first three gradients) is more sensitive than only looking at the first gradient.

## DISCUSSION

The present study aimed to investigate how anaesthesia-induced loss of consciousness alters distributed brain activity in marmosets using fMRI. Expanding upon previous findings from humans and macaques, we investigated brain-wide connectivity patterns through three distinct markers of distributed function; namely, hierarchical structure-function relationships across scales, hierarchical integration and segregation of functional signals, and functional (gradient) geometry. Prior studies in humans and macaques have demonstrated that anesthesia with propofol and sevoflurane leads to increased dependence of functional activity on structural connectivity, as indicated by increased contribution of coarse-grained structural harmonics to the functional signal. In both humans and macaques, reduced gradient range (presumed to indicate diminished depth of the cortical processing hierarchy) was also observed, and in macaques this was accompanied by decreased hierarchical integration of the functional connectome.

In the present study, we examined these markers under propofol and sevoflurane anaesthesia, while also extending our investigation to isoflurane, an anaesthetic not previously examined in this context. In line with previous findings, across all three anaesthetics, we observed a rise in harmonic energy, a contraction of gradient range (with exception of sevoflurane) and overall reduction of gradient dispersion, and decline in hierarchical integration (again with the exception of sevoflurane, which instead exhibited the opposite effect), reinforcing the idea that such alterations in distributed brain-activity are closely related to anaesthesia-induced unconsciousness. Increased energy of connectome harmonics indicates that functional brain activity becomes increasingly constrained by the underlying anatomical white-matter connectivity, reflecting a loss of the flexible, multi-scale interactions seen in the waking state [3]. Anaesthesia concomitantly led to a marked reduction in gradient dispersion [4, 5]. Although a contraction of the principal functional gradient’s range was also observed for two of the three anaesthetics, the dispersion index proved more sensitive, arguably because it takes into account more gradients than just the first.

The present findings are aligned with previous studies investigating the effects of anaesthetics on brain function in marmosets. Previous studies showed decreased functional connectivity under isoflurane [25, 39] and altered resting state network formation in isoflurane and sevoflurane, with less pronounced effects for propofol [24]. In our study, propofol also exhibits the smallest effect size in harmonic energy, gradient dispersion and hierarchical segregation change. However, we still observe significant changes in several measures under the effects of propofol, suggesting increased sensitivity of the measures employed here for detecting the anaesthetic state: possibly because the present approaches take into account the entire functional organisation of the cortex, rather than focusing on discrete cortical patches. On the other hand, hierarchical integration shows the opposite effect for sevoflurane than the other anaesthetics, which represents a point of departure from a previous study in macaques [4], possibly representing a study-specific effect. Although here we have focused on similarities across anaesthetics, future work may also seek to disentangle the unique effects of each, as each of them has a unique profile of receptor affinity.

A pervasive limitation of animal studies is that consciousness cannot be directly addressed via subjective reports, as with humans, requiring reliance on indirect measures such as behavioural responsiveness, which often co-vary with consciousness but are not identical with it and could be dissociated. Future work may seek to validate the present results against additional neural markers of consciousness such as the local-global response or Perturbational Complexity Index, or slow-wave activity saturation of the EEG [40–42]. A limitation of this study is its restricted focus on cortical areas. Given the recent evidence for a prominent role of thalamic nuclei in controlling global states of consciousness [4, 5, 42–44], future work may also focus on this and other subcortical areas - as well as the brainstem, whose importance is also becoming increasingly recognised [45, 46]. Furthermore, the limited number of cortical regions defined by the atlas used constrains the resolution attainable for both functional gradients and harmonic analyses. We also acknowledge that the number of time points in each scan is relatively short. However, this limitation is mitigated by two factors. First, we did not employ time-resolved analyses. Second, short duration was compensated by having several scans for each animal in each condition. Finally, as is typical in similar works, we concentrated on the overall extent of functional gradients rather than their specific topography or other geometric properties inferable from the gradient space. Future studies may benefit from a more comprehensive analysis that considers the broader range of information offered by these methods. Regarding eigenmode computations, we employed a marmoset connectome template, as previously done for both human and macaque; however, use of subject-specific structural connectomes may reveal additional insights in future work. Furthermore, future work may assess whether additional insights can be obtained by using alternative eigenbases, such as obtained from cortical geometry, as suggested by recent work [8] (but see [47] for evidence of the importance of long-range anatomical connections beyond local geometry).

Altogether, the present study demonstrates that anaesthetic-induced loss of consciousness in the marmoset is characterised by a consistent and systematic reorganization of large-scale functional brain architecture, with increased structure-function coupling and a contraction of the hierarchical functional organisation. We have successfully generalised these findings previously observed in humans and macaques to a lissencephalic primate species, and to another widely used anaesthetic agent, isoflurane. Taken together, a consistent pattern emerges: measures of distributed function (gradients, eigenmodes) prove more sensitive to anaesthetic state than local measures, and among them, the most sensitive are those that account for multiple dimensions simultaneously. This consistency bolsters the translational potential of large-scale neuroimaging measures as general measures of the anaesthetic state across species and drugs [2–5]. In turn, consistent findings from marmoset to human underscore the importance of the marmoset as a critical translational bridge in neuroscience, situated between the experimental accessibility and fast developmental timescales of rodents and the complex brain structure of humans and macaques. Ultimately, this work advances our understanding of the fundamental principles governing how the primate brain’s large-scale organization supports the maintenance and loss of consciousness.

## METHODS

### Marmoset FMRI anaesthesia dataset

The marmoset data included in the current study have been published before [24]. For clarity and consistency of reporting, we use the same wording as in the original publication where possible, and we refer the reader to the original work for details.

#### Animals and ethics

This study was approved by the Animal Experiment Committees at the RIKEN Center for Brain Science (CBS) and was conducted per the guidelines for Conducting Animal Experiments of RIKEN CBS. Three male and one female healthy common marmosets (C. jacchus) between 3 and 6 years of age were included. All marmosets were examined 8 times to collect functional MRI data in all conditions. Data was collected in the awake condition first, and sedate/anesthetic data were collected in a random order with an interval of 1 month between each examination in each individual [24].

#### Anesthesia protocol

We used data from awake scans, and from scans under general anaesthesia induced with isoflurane, sevoflurane, or propofol. Essential details are reported below, and we refer to the original publication for additional detail [24].

##### Isoflurane

Three percent isoflurane with 100% *O*_2_ as carrier gas was administered to the marmosets through a facial mask retained with leather gloves. Once sufficient sedation was achieved, an 8 Fr catheter (Atom multi-use tube, Atom Medical Corp., Tokyo, Japan) was inserted into the trachea as an intratracheal tube. The isoflurane concentration was reduced to 2.5%, and 50 *μ*g/kg of atropine and 3.0 mL of physiological saline were administered subcutaneously to prevent intratracheal secretion and dehydration, respectively. After intratracheal intubation, general anesthesia was maintained with 1.8% isoflurane. The marmosets were connected to artificial ventilation for small animals (SN-480-7, Shinano Seisakusho, Tokyo, Japan), and mechanical ventilation was performed under the following conditions: 50% inspiratory oxygen, 8 mL of tidal volume, and 30 breaths per min RR. PR, SpO2, RR, EtCO2, inspiratory and endtidal isoflurane concentrations, and rectal temperature were measured with a vital sign monitor.

##### Sevoflurane

All procedures were performed in the same way as isoflurane, except for dosages. The induction of general anesthesia, intratracheal intubation, and maintenance of general anesthesia were performed with 5.0 and 3.0% of sevoflurane (Pfizer Japan Inc., Tokyo, Japan), respectively.

##### Propofol

Propofol (12 *μ*g/kg) was administered as induction of general anesthesia over 3 min via an indwelling needle. Atropine (50 *μ*g/kg) was administered subcutaneously to prevent intratracheal secretion. Immediately after bolus administration, the predicted plasma concentration of propofol was titrated to 7–9 *μ*g/mL for intratracheal intubation with continuous infusion. The administration dose and protocol were calculated beforehand using a pharmacokinetic parameter reported by [48] and pharmacokinetic analysis software, NONMEM ver. VII (GloboMax ICON Development Solutions, Ellicott City, MD, USA). Once a sufficient plasma concentration was obtained, the propofol dose was controlled to maintain that concentration. Intra-tracheal intubation, respiratory management, and vital sign monitoring were performed in the same manner as for isoflurane.

#### MRI data acquisition

During imaging, the marmosets were placed on a custom-made imaging table (Takashima Seisakusho Co., Ltd, Tokyo, Japan) and immobilized by fixing the head post using a head post fixing tool attached at a custom-made imaging table in all conditions. The marmosets were fitted with earplugs. A hot water circulator was used during imaging to maintain body temperature 36–38 degrees centigrade under all conditions [24]. In Awake condition, data were collected in the dark and monitored with an infrared camera to prevent the marmosets from falling asleep. If the marmosets were observed closing their eyes during a scan, they were awakened with a loud noise before the next scan was started. They were rewarded with a highly palatable food at the end of each imaging. In anesthetic condition, the marmosets were monitored in the same way to observe spontaneous movement or coughing for intratracheal tube. If spontaneous movement or coughing was observed, additional sedatives/anesthetics were administrated and infusion rate or concentration of inhalational anaesthesia was increased.

An ultra-high field MRI system with a static magnetic field strength of 9.4 T (Bruker BioSpin, Ettlingen, Germany), a custom-made 8-channel receiver coil for the marmoset head (Takashima Seisakusho Co., Ltd, Tokyo, Japan), and a 154 mm inner diameter transmitter coil (Bruker BioSpin, Ettlingen, Germany) were used to collect structural and functional data. Structural data and T2-weighted images were imaged using rapid acquisition with relaxation enhancement (RARE) sequence with the following conditions and parameters: time repetition (TR)=4331 ms, time echo (TE) = 15.0 ms, FOV = 42.0 × 28.0 × 36.0 mm, matrix size = 120 × 80 voxels, resolution = 0.35 × 0.35 mm, slice thickness = 0.7 mm, number of slices = 52, scan time = 1 min and 26 s, RARE factor = 4. Functional images were captured using a gradient recalled echo-planar imaging (EPI) sequence with the following conditions and parameters: TR = 2,000 ms, TE = 16.0 mm, FOV = 42.0 × 28.0 × 36.0,mm matrix size = 60 × 40 voxels, resolution = 0.7 × 0.7 mm, slice thickness = 0.7 mm, number of slices = 52, repetition = 155, scan time = 310s. Functional imaging was performed 12 times per animal, per condition [24].

#### Marmoset functional MRI preprocessing and denoising

The preprocessing procedure for the dataset utilised have been previously reported in [24]. Essential details are reported, and we refer to the original publication for additional detail

After the acquired data were converted to Neuro Informatics Technology Initiative format (NIfTI), the voxel size was changed from 0.7 mm isotropic to 3.5 mm isotropic using SPM (Wellcome Trust Center for Neuroimaging, London, UK). the top-up tool of the FM-RIB Software Library (FSL) software (FMRIB, Oxford, UK) was utilised for the estimation and correction of geometric distortions induced by magnetic susceptibility, as a single excitation in EPI was used for imaging of all cross-sections. Discrepancies in signal acquisition timing across sections were corrected using slice time correction. Head movements caused by body movements was corrected using realignment. Six directions of deviation were accounted for: x (left/right), y (front/back), z (up/down), pitch (rotational direction of nodding and looking up), roll (rotational direction of moving the ear closer to the shoulder), and yaw (rotational direction of looking left/right). For each time repetition (TR), the deviation from the reference time point, and the initial functional brain image, was determined; and the image was moved and rotated by the rigid body model based on this deviation. The method of finding the parameters of the linear transformation was used to minimize the difference between the first functional brain image and the affine transformation of the series of functional brain images to be corrected, by calculating convergence using the method of least squares. After correcting the spatial scale error between the structural and functional images with co-registration, segmentation was performed to provide information on the tissue to which each voxel belongs in terms of brain tissue classification. The voxels spatial standardisation through normalisation was executed, aligning voxels to the standard brain image, therein correcting for structural differences between marmosets. Excessive voxel value fluctuations within individuals were suppressed by smoothing and application of normal probability field theory. Functional data were smoothed using spatial convolution with a Gaussian kernel of 2 voxels (7 mm). Ordinary least squares regression with cerebrospinal fluid pulsation, heart rate, and respiratory artifacts as regressors was used for denoising of physiological noise. Temporal band pass filtering was performed by frequency filtering (0.01–0.1 Hz) using the fMRI denoising pipeline of CONN toolbox. Finally, preprocessed functional data were parcellated into 70 regions in the cerebral cortex, corresponding to regions of the marmoset MBM atlas [49] for which structural connectivity data was also available (see below).

#### Marmoset structural connectome

The marmoset structural connectome was obtained from the Brain/MINDS Marmoset MRI NA216 in-vivo database (https://doi.org/10.24475/bminds.mri.thj.4624) [50]. Full details are provided in the original data-release publication [50]. Briefly: in vivo MR imaging was conducted using a 9.4 T BioSpec 94/30 (Bruker Optik GmbH, Ettlingen, Germany) unit and a transmitting and receiving coil with an 86-mm inner diameter. Diffusion weighted spin-echo echo planar imaging was used with TR = 3000 ms, TE = 25.6 ms, b-value = 1000 and 3000 s/mm2 in 30 and 60 diffusion directions, respectively (plus 4 b0 images), number of segments = 6, FA = 90, NA = 3, voxel size = 350 × 350 × 700 micrometers, and scan time = 90 min. Diffusion metrics were created by DTI model, and diffusion fiber structural connectome was created by constrained spherical deconvolution, using the number of streamlines between each pair of 104 brain regions (37 cortical and 15 subcortical per hemisphere) [50]. Here we used the structural connectivity between 70 pairs of regions (35 per hemisphere) which correspond to the cortical regions of the marmoset MBM atlas [49].

### Analysis

The analysis techniques utilised in the current study were previously reported in [4]. Essential details are outlined below, and we refer to the original publication for additional details.

#### Structure-function coupling via harmonic mode decomposition

Functional spatiotemporal patterns of neural activity derived from fMRI can be decomposed in terms of anatomically-based distributed building blocks: eigen-vectors of the graph Laplacian of the structural connectome [3, 12, 13]. Here, as in previous work [4] we draw inspiration from the original Connectome Harmonic

Decomposition framework outlined by Atasoy and colleagues [12, 13], who used the harmonic modes of a high-resolution human structural connectome, obtained from combining long-range white matter tracts and local connectivity within the grey matter. It is important to note that here we use harmonic modes derived from a parcellated connectome of diffusion MRI and tract-tracing, and therefore refer to the eigenmodes obtained in this way as “harmonic modes”, preserving the original nomenclature, associating the term “connectome harmonics” with the harmonic modes originally developed by Atasoy and colleagues.

Following the method developed by Atasoy and colleagues [12, 13], we compute the symmetric graph Laplacian Δ_*G*_ on the matrix *C* which represents the marmoset structural connectome. To ensure symmetry of the corresponding connectivity matrix, aquisition of real eigenvalues and obtaining an undirected connectome, we average entries above and below the diagonal. From here, we estimate the connectome Laplacian (the discrete counterpart of the Laplace operator Δ applied to the network of the marmoset structural brain connectivity):

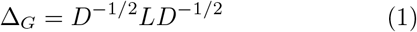

with *L* = *D* − *C*, where *D* is the diagonal “degree matrix” of the graph *C* i.e.

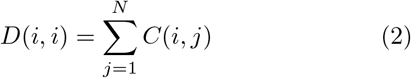

We then calculate the harmonic modes *φ*_*k*_, *k* ∈ {1, …, *N*} by solving the following eigenvalue equation:

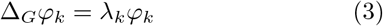

∀*k* ∈{1,…, *N*, with 0 *< λ*_1_ *< λ*_2_ *<* … *< λ*_*n*_ where *λ*_*k*_, *k* ∈{1, …, *N*} is the corresponding eigenvalue of the eigenvector *φ*_*k*_ . In other words, *λ*_*k*_ and *φ*_*k*_ are the eigen-values and eigenvectors of the Laplacian of the primate structural connectivity matrix (marmoset connectome). Therefore, if *φ*_*k*_ is the harmonic pattern of the *k*^*th*^ spatial frequency, then the corresponding eigenvalue is a term relating to the intrinsic energy of that particular harmonic mode (see figure 1). With an increasing harmonic number *k*, we obtain more complex and fine-grained spatial patterns (see Figures 5 and 6).

#### Decomposition of fMRI data

At each timepoint *k* ∈ {1, …, *T* }, (corresponding to one TR), the spatial pattern of cortical activity over brain regions at time t, denoted as Ft, was decomposed as a linear combination of the set of harmonic modes 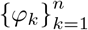:

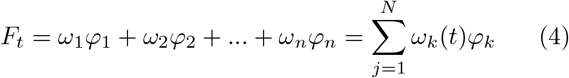

with the contribution *ω*_*k*_(*t*) of each harmonic mode *φ*_*k*_(*t*) at time *t* being estimated as the projection (dot product) of the fMRI data *F*_*t*_ onto *φ*_*k*_:

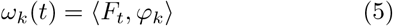

#### Energy of harmonic modes

Once the fMRI cortical activation pattern at time has been decomposed into a linear combination of harmonic modes, the magnitude of each harmonic’s contribution to the cortical activity of each harmonic *φ*_*k*_, *k* ∈ {1, …, *n*}, (regardless of sign) at any given timepoint *t*, denoted *P* (*φ*_*k*_, *t*), is called its “power”, for analogy with the Fourier transform, is computed as the amplitude of its contribution:

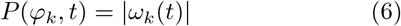

The normalized frequency-specific contribution of each harmonic *φ*_*k*_, *k* ∈{1, …, *n*} at timepoint *t*, termed “energy”, is estimated by combining the magnitude strength of activation (power) of a particular harmonic mode with its own intrinsic energy given by the associated eigenvalue 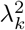:

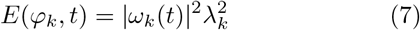

#### Functional Gradient Mapping from Diffusion Map Embedding

Marmoset cortical functional gradients were calculated using the BrainSpace toolbox https://github.com/MICA-MNI/BrainSpace [7] as implemented in MATLAB, with default parameters for density, similarity kernel, and anisotropic diffusion parameter (see below).

The functional connectivity matrix (FC) is calculated as the Pearson correlation between each pair of regional fMRI signals per scan per condition.

Following previous work, each matrix was z-transformed and thresholded row-wise to achieve 10% density (i.e., retaining 10% of entries), retaining only the strongest connections in each row [4, 7]. The normalised cosine similarity was calculated on the thresholded z-matrix to generate a similarity matrix reflecting the similarity in whole-brain connectivity patterns between vertices. While the FC matrix reflects how similar each pair of regions are in terms of their temporal co-fluctuations, this similarity matrix reflects the similarity of a pair of regions with respect to their patterns of FC. The similarity matrix is required as input to the diffusion map embedding algorithm that we have used here in agreement with previous work on functional gradients and how they are reshaped by pharmacological interventions [7].

Diffusion Map Embedding is a nonlinear manifold learning technique [6, 51] that exploits the properties of the graph Laplacian to model the diffusion process, and is therefore related to harmonic mode decomposition - though performed on functional data rather than using a common structural connectome, to reveal its contributions to the functional activations (indeed, a deep mathematical analogy exists between diffusion graph embedding on the functional connectome and the recent extension of CHD called “functional harmonics” [52]).

A further parameter (*α*) controls the relative influence of sampling points density on the manifold. The parameter sits in the range of 0 to 1, and in the case of diffusion map embedding is set to 0.5 to provide a balance between local and global contributions to the embedding space estimation [7].

The high-dimensional similarity matrix is treated as a graph, with connections (entries of the similarity matrix) reflecting the similarity between the regional FC patterns. The technique estimates a low-dimensional set of embedding components (gradients); in this low-dimensional space, proximity reflects similarity of the patterns of FC: regions with greater similarity in FC patterns (those more strongly connected in the network) are placed are more proximal to one another, and regions with lower similarity are placed further apart. In this way, each gradient represents one dimension of covariance in the inter-regional similarity between FC patterns, with a small number of gradients capturing most of the dimensions of inter-regional similarity, this can then be visualised through a low-dimensional scatter plot [6, 7]. In the embedding space, each gradient can be understood to be “anchored” at regions that have the strongest values for that gradient, suggesting that this particular embedding dimension captures their respective similarity profiles well. In contrast, regions that are close to the origin (i.e. have a low absolute value for a particular gradient) mean that they are only minimally similar to the “anchor points” of that gradient, which thereby does not strongly capture their overall FC similarity profile well [6, 7].

Therefore, the greater the difference between the extremes of a gradient, the more the differentiation between regions is being captured by that gradient. To quantify this formally, we calculated the difference between the maximum and minimum values of each scan along the first gradient [7, 11] (which mathematically captures most of the variation in FC profiles within each scan) and compared these differences across conditions. Note that the dimension of greatest variability (first gradient) is assessed in a data-driven manner and need not be identical across different scans. The range of the second gradient is computed in the same way. Additionally, we consider the dispersion of the first three gradients, calculated as per Huang et al [5] as the sum of squared Euclidean distance of all regions to the global centroid in the 3D cortical gradient space. Finally, we also consider an additional measure, the ratio of the eigenvalue associated with the first gradient, to the sum of all gradients’ eigenvalues, reflecting its relative importance.

To demonstrate the robustness of our results, we show that they are replicated even when the parameter is set to very different values (0.1 or 0.9); when using lower sparsity levels for the input FC matrix (50% sparsity or 10% sparsity); and when using cosine similarity instead of normalised angle similarity (see supplementary figures S2-S5 and S7-S10).

#### Hierarchical Integration and Segregation

A third perspective on distributed brain function that can be obtained from studying the brain’s eigenmodes emerges from its hierarchical organisation, and how the latter supports the balance between integration and segregation across scales - which is a central feature of many prominent scientific theories of consciousness. Classical graph-theoretic measures quantify integration and segregation at a one scale and are thus not suitable for capturing these properties across multiple hierarchical modules [36, 38]. However, a recently introduced formalism based on eigenmodes of functional connectivity can provide such quantification [17].

The functional connectivity (*FC*) is a symmetric matrix, which can be decomposed as *FC* = *U* Λ*U*^*T*^ where *U* is an orthogonal matrix whose columns are eigenvectors (eigenmodes) of *FC*, and Λ is a diagonal matrix whose entries are the eigenvalues of *FC*. Each FC eigenmode identifies a distinct pattern of regions that are coactivated (same sign) or alternately activated (opposite sign). Therefore, hierarchical modules can be identified based on the concordance or discordance of signs between regions across eigenmodes, progressively partitioning the *FC* into a more and progressively smaller modules: starting with the primary eigenmode with all regions exhibiting the same sign, up to the level where each module coincides with a single region, indicative of completely segregated activity. Thus, segregated modules at one level of the hierarchy can become integrated by being part of the same superordinate module. During this nested partitioning process, we obtain the module number *M*_*i*_, (*i* =1… *N* ) and the modular size *M*_*j*_, (*j* = 1… *N* ) at each level.

Each level *i* of the hierarchy is characterised by two quantities: the number of modules *M*_*i*_ into which the cortex is divided, and the covariance explained by the corresponding eigenmode (given by its squared eigenvalue 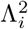 ). However, the number of modules alone may not properly describe the picture of nested segregation and integration due to heterogeneity in module sizes. The correction factor was calculated as *p*_*i*_ = Σ_*j*_ |*m*_*j*_ *− N/M*_*i*_|*/N* which reflects the deviation from the optimised modular size in the *i*^*th*^ level. Thus, the correction effect is stronger for a larger deviation of modular size from homogeneity [17].

Since the first eigenmode encompasses the entire cortex into one whole-brain module, the corresponding eigen-value 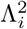 quantifies the overall contribution of whole-brain integration,

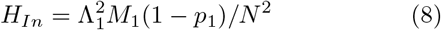

The overall level of segregation across the hierarchy is quantified as

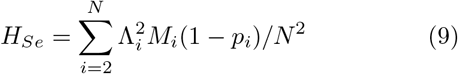

the sum of contributions from all eigenmodes except the first (i.e., all eigenmodes that involve a partitioning of the cortex), each weighted by the corresponding number of modules *M*_*i*_ (further corrected for heterogeneous modular sizes, since a partition into two modules of size 1 and *N −* 1 is clearly not as segregated as a partition into two equally-sized modules).

### Statistical Reporting

Significance was assessed using linear mixed effects modelling (implemented using MATLAB’s *fitlme* function), with condition as the fixed effect and animal identity as the random effect. This approach enabled us to take into account the fact that the same animal could provide more than one data-point to each condition, as well as contributing data-points for more than one condition. Beta coefficients are reported, alongside 95% confidence intervals (lower, upper). P-values are corrected for comparisons across multiple anaesthetic conditions using the False Discovery Rate procedure.

## Data availability

The marmoset fMRI data are available from author KM for scientific collaboration. The marmoset structural connectome is available from the Brain/MINDS Marmoset MRI NA216 in-vivo database (https://doi.org/10.24475/bminds.mri.thj.4624).

## Code availability

The BrainSpace toolbox is freely available online at https://github.com/MICA-MNI/BrainSpace. MATLAB code from [17] to quantify hierarchical integration and segregation is freely available online at https://github.com/TobousRong/Hierarchical-module-analysis.

## ACKNOWLEDGEMENTS

AIL and HA are supported by a Wellcome Early-Career Award (grant number 226924/Z/23/Z). DO was supported by a FENS/IBRO-PERC Exchange Fellowship. KJ is supported by the Centre for Eudaimonia and Human Flourishing, Linacre College, University of Oxford (funded by the Pettit and Carlsberg Foundations). ES is supported by the Luxembourg National Research Fund (FNR) (17906488). This work was also supported by the program for Brain Mapping by Integrated Neurotechnologies for Disease Studies (Brain/MINDS) from the Japan Agency for Medical Research and Development (AMED) (Grant Number JP23wm0625001 to HO), JSPS KAKENHI (Grant Number JP20H03630 to JH), and by “MRI platform” as a program of Project for Promoting public Utilization of Advanced Research Infrastructure of the Ministry of Education, Culture, Sports, Science and Technology (MEXT), Japan (Grant Number JPMXS0450400622 to JH). M.L.K. is supported by the Center for Music in the Brain, funded by the Danish National Research Foundation (DNRF117), and Centre for Eudaimonia and Human Flourishing at Linacre College funded by the Pettit and Carlsberg Foundations. G.D. is supported by grant no. PID2022-136216NB-I00 funded by MICIU/AEI/10.13039/501100011033 and by ‘ERDF A way of making Europe’, ERDF, EU, Project NEurological MEchanismS of Injury, and Sleep-like cellular dynamics (NEMESIS; ref. 101071900) funded by the EU ERC Synergy Horizon Europe, and AGAUR research support grant (ref. 2021 SGR 00917) funded by the Department of Research and Universities of the Generalitat of Catalunya. For the purpose of open access, the authors have applied a Creative Commons Attribution (CC BY) licence to any Author Accepted Manuscript version arising from this submission.

