## Supplementary Information for "Generalisable signatures of anaesthesia in the large-scale functional organisation of the marmoset brain"

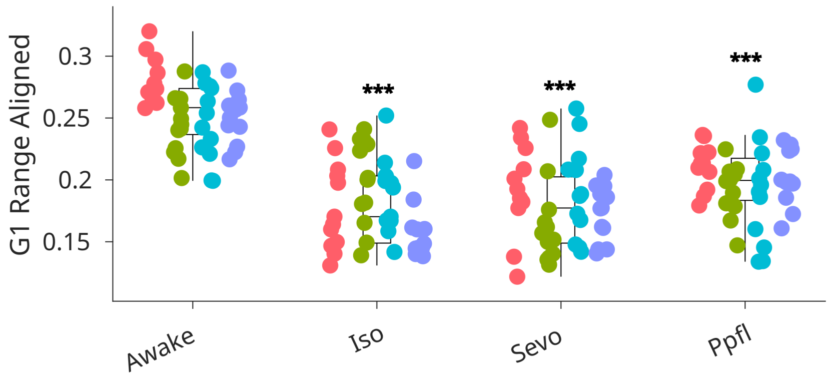


Figure S1. **Effect of three distinct anaesthetics on the range of the aligned principal functional gradient, derived from the group average functional connectivity from the Awake condition.** $*, p < 0.05$; $**, p < 0.01$; $* * *, p < 0.001$ (derived from linear mixed effects modelling; two-sided, FDR corrected) anaesthesia conditions compared against Awake condition. Box plots: central line denotes median; box limits signify the upper and lower quartiles; whiskers signify 1.5× interquartile range. Within each condition (defined on X axis) values of the same colour are measured from the same marmoset.

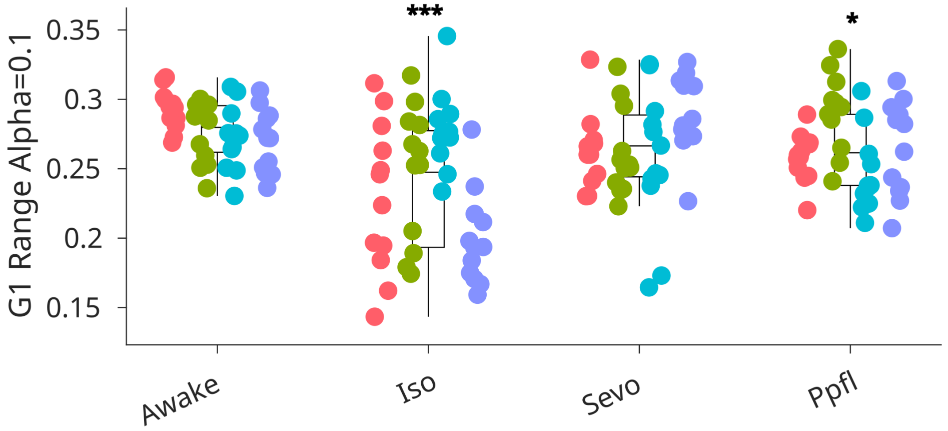


Figure S2. **Effect of three distinct anaesthetics on the range of the principal functional gradient with the diffusion parameter $\alpha$ set to 0.1.** $*$, $p < 0.05$; $**$, $p < 0.01$; $***$, $p < 0.001$ (derived from linear mixed effects modelling; two-sided, FDR corrected) anaesthesia conditions compared against Awake condition. Box plots: central line denotes median; box limits signify the upper and lower quartiles; whiskers signify 1.5× interquartile range. Within each condition (defined on X axis) values of the same colour are measured from the same marmoset.

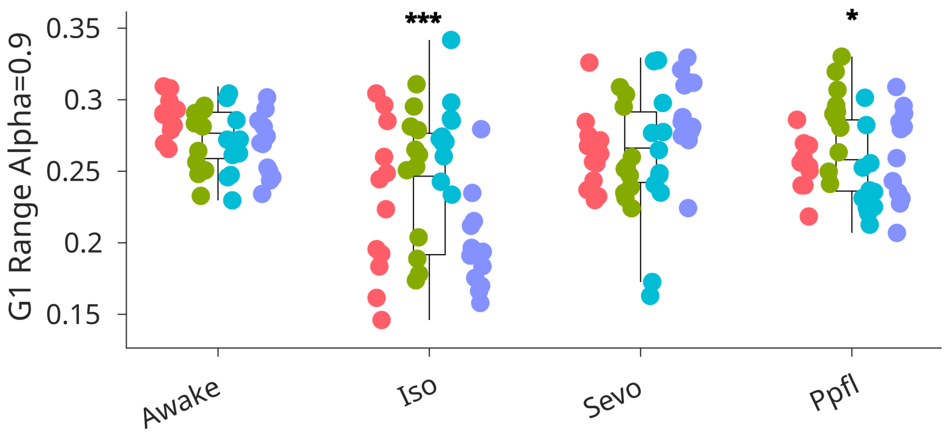


Figure S3. **Effect of three distinct anaesthetics on the range of the principal functional gradient with the diffusion parameter $\alpha$ set to 0.9.** $*$, $p < 0.05$; $**$, $p < 0.01$; $***$, $p < 0.001$ (derived from linear mixed effects modelling; two-sided, FDR corrected) anaesthesia conditions compared against Awake condition. Box plots: central line denotes median; box limits signify the upper and lower quartiles; whiskers signify 1.5$\times$ interquartile range. Within each condition (defined on X axis) values of the same colour are measured from the same marmoset.

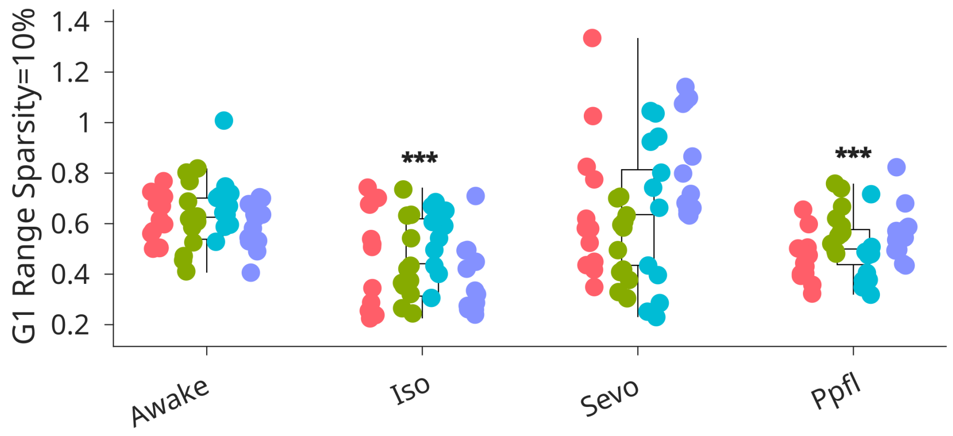


Figure S4. **Effect of three distinct anaesthetics on the range of the principal functional gradient range with the sparsity parameter set to 10%.** $*$, $p < 0.05$; $**$, $p < 0.01$; $***$, $p < 0.001$ (derived from linear mixed effects modelling; two-sided, FDR corrected) anaesthesia conditions compared against Awake condition. Box plots: central line denotes median; box limits signify the upper and lower quartiles; whiskers signify 1.5$\times$ interquartile range. Within each condition (defined on X axis) values of the same colour are measured from the same marmoset.

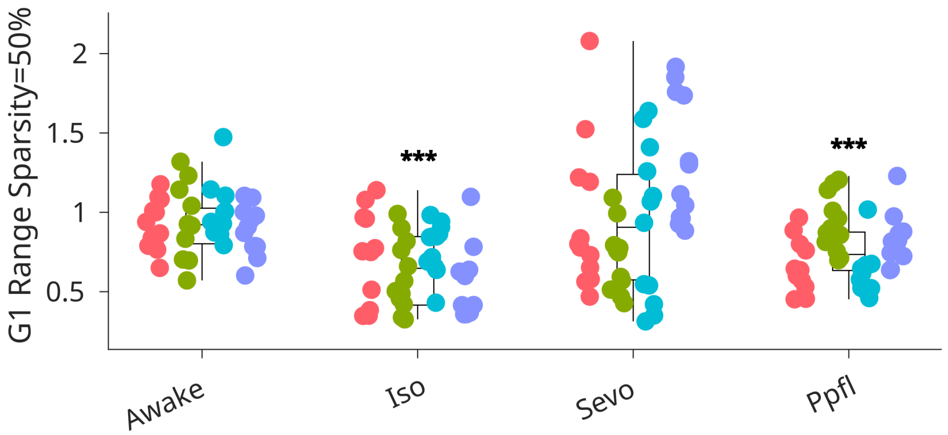


Figure S5. **Effect of three distinct anaesthetics on the range of the principal functional gradient range with the sparsity parameter set to 50%.** $*$, $p < 0.05$; $**$, $p < 0.01$; $***$, $p < 0.001$ (derived from linear mixed effects modelling; two-sided, FDR corrected) anaesthesia conditions compared against Awake condition. Box plots: central line denotes median; box limits signify the upper and lower quartiles; whiskers signify 1.5× interquartile range. Within each condition (defined on X axis) values of the same colour are measured from the same marmoset.

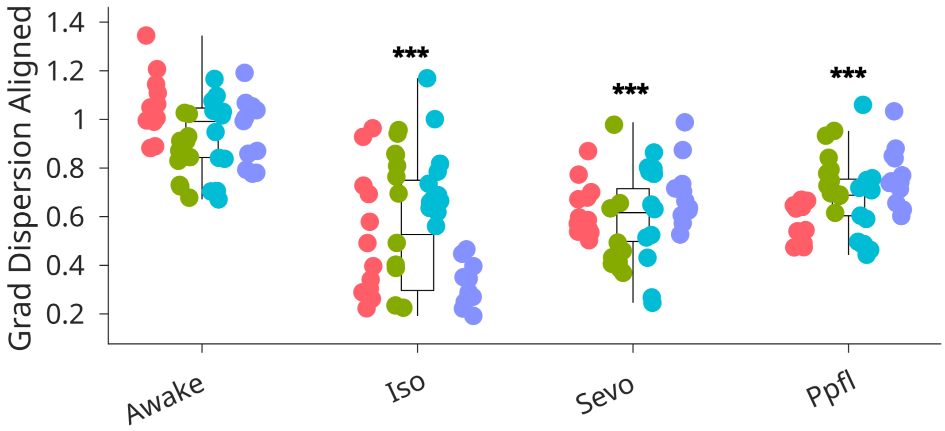


Figure S6. **Effect of three distinct anaesthetics on the dispersion of the first three aligned gradients, each derived from the group average functional connectivity from the Awake condition**. $*$, $p < 0.05$; $**$, $p < 0.01$; $***$, $p < 0.001$ (derived from linear mixed effects modelling; two-sided, FDR corrected) anaesthesia conditions compared against Awake condition. Box plots: central line denotes median; box limits signify the upper and lower quartiles; whiskers signify 1.5× interquartile range. Within each condition (defined on X axis) values of the same colour are measured from the same marmoset.

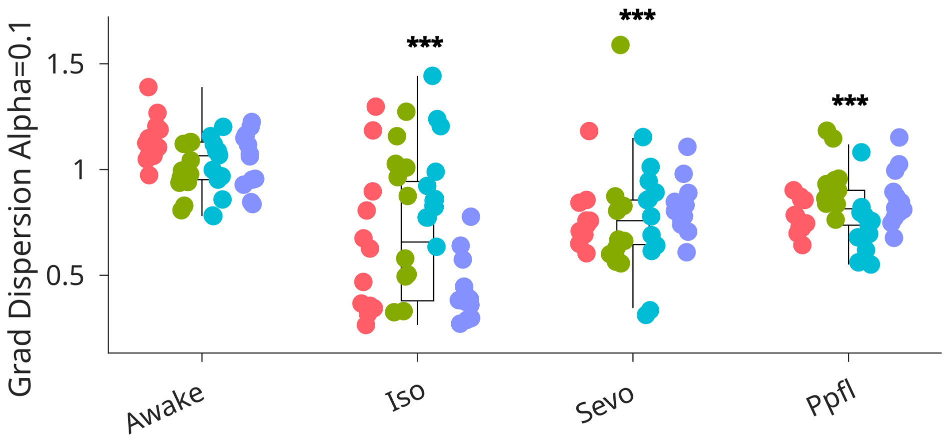


Figure S7. **Effect of three distinct anaesthetics on the dispersion of the first three gradients with the diffusion parameter $\alpha$ set to 0.1.** $*$, $p < 0.05$; $**$, $p < 0.01$; $***$, $p < 0.001$ (derived from linear mixed effects modelling; two-sided, FDR corrected) anaesthesia conditions compared against Awake condition. Box plots: central line denotes median; box limits signify the upper and lower quartiles; whiskers signify 1.5$\times$ interquartile range. Within each condition (defined on X axis) values of the same colour are measured from the same marmoset.

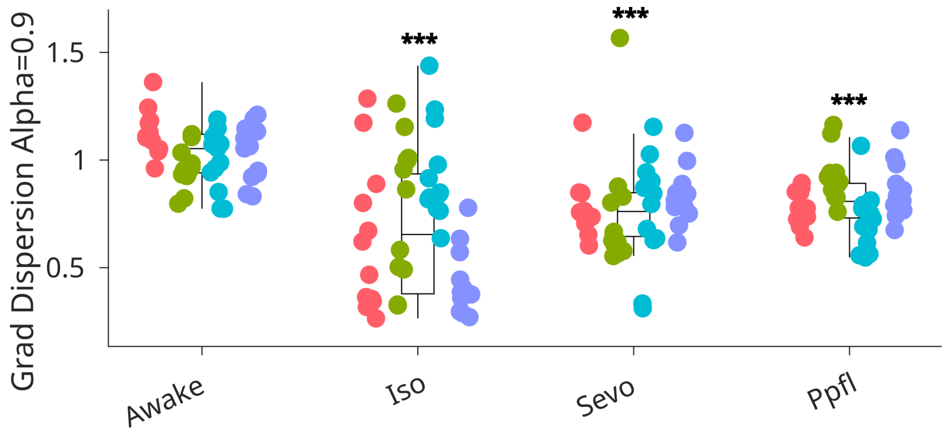


Figure S8. **Effect of three distinct anaesthetics on the dispersion of the first three gradients with the diffusion parameter $\alpha$ set to 0.9.** $*$, $p < 0.05$; $**$, $p < 0.01$; $***$, $p < 0.001$ (derived from linear mixed effects modelling; two-sided, FDR corrected) anaesthesia conditions compared against Awake condition. Box plots: central line denotes median; box limits signify the upper and lower quartiles; whiskers signify 1.5$\times$ interquartile range. Within each condition (defined on X axis) values of the same colour are measured from the same marmoset.

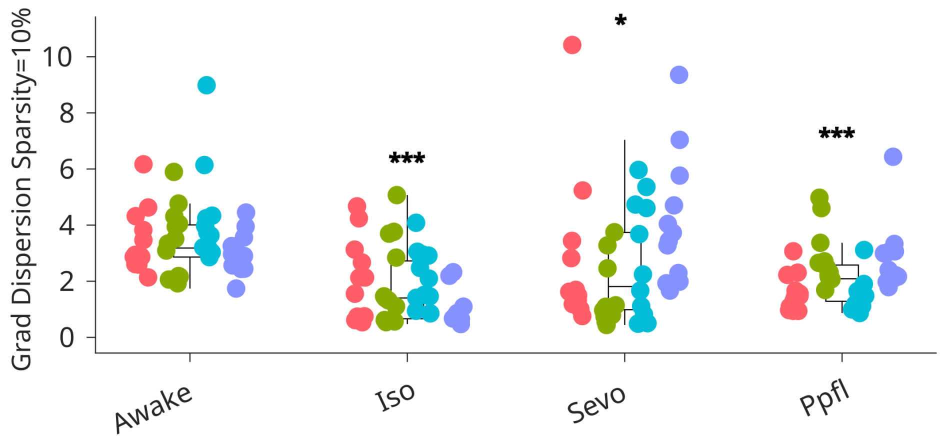


Figure S9. **Effect of three distinct anaesthetics on the dispersion of the first three gradients with the sparsity parameter set to 10%.** $*$, $p < 0.05$; $**$, $p < 0.01$; $***$, $p < 0.001$ (derived from linear mixed effects modelling; two-sided, FDR corrected) anaesthesia conditions compared against Awake condition. Box plots: central line denotes median; box limits signify the upper and lower quartiles; whiskers signify 1.5× interquartile range. Within each condition (defined on X axis) values of the same colour are measured from the same marmoset.

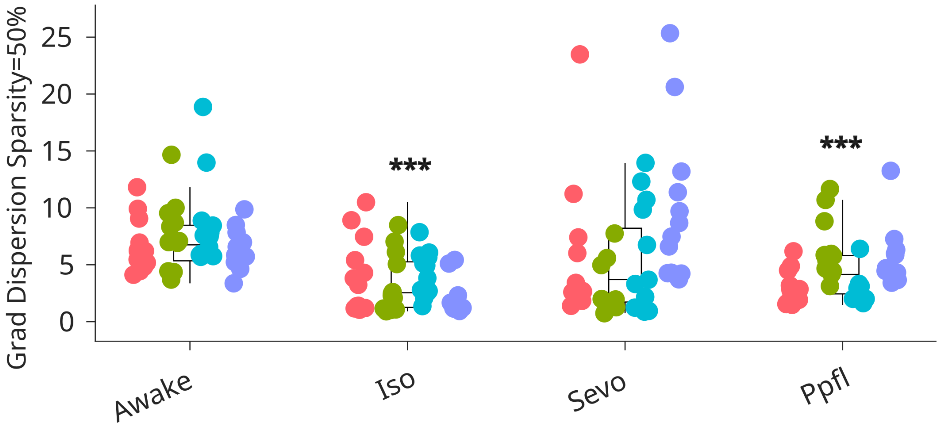


Figure S10. **Effect of three distinct anaesthetics on the dispersion of the first three gradients with the sparsity parameter set to 50%.** $*$, $p < 0.05$; $**$, $p < 0.01$; $***$, $p < 0.001$ (derived from linear mixed effects modelling; two-sided, FDR corrected) anaesthesia conditions compared against Awake condition. Box plots: central line denotes median; box limits signify the upper and lower quartiles; whiskers signify 1.5× interquartile range. Within each condition (defined on X axis) values of the same colour are measured from the same marmoset.

| Measure | Contrast | Estimate | SE | Lower | Upper | DF | tStat | p_FDR | Sig |
|---|---|---|---|---|---|---|---|---|---|
| Harmonic Energy | Awake vs iso | 6.174 | 0.573 | 5.036 | 7.312 | 94 | 10.771 | <0.001 | *** |
| Harmonic Energy | Awake vs propo | 2.394 | 0.418 | 1.564 | 3.225 | 94 | 5.726 | <0.001 | *** |
| Harmonic Energy | Awake vs sevo | 3.194 | 0.485 | 2.23 | 4.158 | 94 | 6.579 | <0.001 | *** |

Table S1. Harmonic Energy

| Measure | Contrast | Estimate | SE | Lower | Upper | DF | tStat | p_FDR | Sig |
|---|---|---|---|---|---|---|---|---|---|
| Gradient Range 1 | Awake vs iso | -0.038 | 0.007 | -0.053 | -0.024 | 94 | -5.271 | <0.001 | *** |
| Gradient Range 1 | Awake vs propo | -0.013 | 0.005 | -0.023 | -0.002 | 94 | -2.382 | 0.029 | * |
| Gradient Range 1 | Awake vs sevo | -0.009 | 0.006 | -0.021 | 0.003 | 94 | -1.469 | 0.145 | n.s. |

Table S2. Gradient Range 1

| Measure | Contrast | Estimate | SE | Lower | Upper | DF | tStat | p_FDR | Sig |
|---|---|---|---|---|---|---|---|---|---|
| Grad Dispersion | Awake vs iso | -0.338 | 0.051 | -0.438 | -0.237 | 94 | -6.663 | <0.001 | *** |
| Grad Dispersion | Awake vs propo | -0.216 | 0.028 | -0.272 | -0.161 | 94 | -7.744 | <0.001 | *** |
| Grad Dispersion | Awake vs sevo | -0.26 | 0.035 | -0.331 | -0.19 | 94 | -7.345 | <0.001 | *** |

Table S3. Grad Dispersion

| Measure | Contrast | Estimate | SE | Lower | Upper | DF | tStat | p_FDR | Sig |
|---|---|---|---|---|---|---|---|---|---|
| Hierarchical Int | Awake vs iso | -0.01 | 0.002 | -0.014 | -0.005 | 94 | -4.6 | <0.001 | *** |
| Hierarchical Int | Awake vs propo | -0.004 | 0.002 | -0.008 | -0.001 | 94 | -2.422 | 0.017 | * |
| Hierarchical Int | Awake vs sevo | 0.015 | 0.004 | 0.008 | 0.022 | 94 | 4.178 | <0.001 | *** |

Table S4. Hierarchical Integration

| Measure | Contrast | Estimate | SE | Lower | Upper | DF | tStat | p_FDR | Sig |
|---|---|---|---|---|---|---|---|---|---|
| Hierarchical Seg | Awake vs iso | -0.024 | 0.004 | -0.032 | -0.017 | 94 | -6.599 | <0.001 | *** |
| Hierarchical Seg | Awake vs propo | -0.024 | 0.003 | -0.03 | -0.017 | 94 | -7.494 | <0.001 | *** |
| Hierarchical Seg | Awake vs sevo | -0.053 | 0.004 | -0.06 | -0.046 | 94 | -15.14 | <0.001 | *** |

Table S5. Hierarchical Segregation
